# Karyotype Description and Comparative Chromosomal Mapping of 5S rDNA in 21 species

**DOI:** 10.1101/2024.02.08.579556

**Authors:** Xiaomei Luo, Yunke Liu, Xiao Gong, Meng Ye, Qiangang Xiao, Zhen Zeng

## Abstract

5S rDNA is essential component of all cell types. A survey was conducted to study the 5S rDNA site number, position, and origin of signal pattern diversity in 64 plants using fluorescence *in situ* hybridization. The species used for this experiment were chosen due to the discovery of karyotype rearrangement, and for species checked in which 5S rDNA has not yet been explored. The chromosome number was 14–160, while the chromosome length was 0.63–6.88 μm, with 37 plants (58%) had small chromosome (<3 μm). The chromosome numbers of three species and 5S rDNA loci of 19 species have been reported for the first time. 5S rDNA was varied and abundant in signal site number (2–18), position (e.g., interstitial, distal, and proximal position, occasionally, outside chromosome), and even as intense (e.g., strong, weak, and slight). Our results exposed modifiability in the number and location of 5S rDNA in 33 plants and demonstrated an extensive stable number and location of 5S rDNA in 31 plants. The potential origin of signal pattern diversity was probably caused by chromosome rearrangement (e.g., deletion, duplication, inversion, translocation), polyploidization, self-incompatibility, and chromosome satellites. These data characterized the variability of 5S rDNA within the karyotypes of the 64 plants that exhibited massive chromosomal rearrangements and provided anchor points for genetic physical maps. These data prove the utility of the 5S rDNA oligonucleotide as a chromosome marker in identifying plant chromosomes. This method provides a basis for developing similar applications for cytogenetic analysis in other species.

**Article Summary:** A survey was exploited to study 5S rDNA signal pattern diversity in 64 plants by FISH. The chromosome number of three species and 5S rDNA loci of 19 species have been reported by the first time. 5S rDNA was really rather various and abundant in signal site number (2–18), position (interstitial, distal, proximal position, occasionally, outside chromosome), even as the intense (strong, week, slight). These data prove the utility of 5S rDNA oligonucleotide as chromosome marker in identifying plant chromosomes.

## Introduction

Ribosomal DNA (rDNA) is essential to all cell types that code rRNA. rDNAs are generally detected by exercising considerable parts of chromosomes, including both 45S and 5S rDNA (Slidler 2016). The 5S rDNA structured numerous copies in genomes and has been far and widely used in hundreds of cytogenetic investigations as an important marker for fluorescence *in situ* hybridization (FISH) analysis (Heslop-Harrison 2000, Rodríguez-González et al. 2023), not only for essential crops (Kamisugi et al. 1994, Luo et al. 2018a, b, c), but also for woody plants, such as walnut (Luo and Chen 2020), the Chinese pepper (Luo et al. 2018d, He et al. 2023, Hu et al. 2023), and wintersweet (Luo and Chen 2019).

The lengths of the 5S rDNA sequence ranged from 48 bp (Turner et al. 2005) to 854 bp (Liu et al. 2017) because of its high copies and variations when searching for “5S rDNA” in the database Nucleotide of National Center for Biotechnology Information. Furthermore, the lengths of 5S rDNA as a FISH probe in the PubMed database from NCBI varied considerably: 41 bp (Luo et al. 2017, Islam-Faridi et al. 2020), 94 bp (Sergeeva et al. 2017), 117 bp (Lukjanová et al. 2023), 120 bp (Taketa et al. 2001, Symonová et al. 2017, Deon et al. 2022, de Moraes et al. 2023), 124 bp (Waminal et al. 2018), 131 bp (Sergeeva et al. 2017), 222 bp (Glugoski et al. 2018), 285 bp (Röser et al. 2001), 300 bp (Araya-Jaime et al. 2022), 302 bp (Zhang et al.2016), 303 bp (Mahelka et al. 2013), 320 bp (Khensuwan et al. 2023), 324 bp (Kamisugi et al. 1994), 326 bp (Robledo and Seijo 2008), 329 bp (Röser et al. 2001), 347 bp, ∼400 bp (Pedrosa et al. 2002), 410 bp (Kovács et al. 2023), 456 bp (Röser et al. 2001), 468 bp, 473 bp, 477 bp, 496 bp (Martins et al. 2000), 497 bp (Alexandrov et al. 2022), 498 bp (Martins et al. 2000), 500 bp (de Barros et al. 2023), 556 bp (Gottlob-McHugh et al. 1990), 596 bp (Joshi et al. 2023), 702 bp (Glugoski et al. 2018), 871 bp (Amarasinghe and Carlson 1988), and 1193 bp (Glugoski et al. 2020).

Conversely, 5S rDNA FISH signal sites in previous research ranged from 1 to 71 (Rodríguez-González et al. 2023). Quite of species conserve 5S rDNA, including only two stable chromosomes with two 5S rDNA signal sites (Zhang et al. 2016, Moraes et al. 2022, Yurkevich et al. 2022, Alexandrov et al. 2022, Khensuwan et al. 2023). Nevertheless, a few species occupy visible 5S rDNA FISH signals, including the numbers of both major and visible dispersed sites. For example, four 5S rDNA signal sites in six in *Sinapidendron frutescens* (Aiton) Lowe (Ali et al. 2005), eight in *Prunus pseudocerasus* (Lindl.) G. Don (Wang et al. 2022), 8-19 in *Trifolium medium* L. (Lukjanová et al. 2023), 10 in *Brassica juncea* (Linnaeus) Czernajew, 12 in *Olimarabidopsis cabulica* (J. D. Hooker & Thomson) Al-Shehbaz et al., 14 in *Brassica napus* L. (Ali et al. 2005), 15 in *Paphiopedilum sukhakulii* Schoser & Senghas (Lan and Albert 2011), 16 in *Piptanthus concolor* Harrow ex Craib (Luo et al. 2017), 19 in *Paphiopedilum henryanum* Braem, 20 in *Paphiopedilum druryi* (Bedd.) Stein, 22 in *Paphiopedilum sangii* Braem, 23 in *Paphiopedilum tigrinum* Koop. et Haseg., 26 in *Paphiopedilum liemianum* Fowlie, 27 in *Paphiopedilum hirsutissimum* (Lindl. ex Hook.) Stein, 28 in *Paphiopedilum victoria-regina* (Sander) M.W.Wood, 29 in *Paphiopedilum primulinum* M. W. Wood et P. Taylor, 30 in *Paphiopedilum glanduliferum* (Blume) Stein, 32 in *Paphiopedilum adductum* Asher, 34 in *Paphiopedilum randsii* Fowlie, 36 in *Paphiopedilum parishii* (Rchb. F.) Stein, and 38 in *Paphiopedilum gigantifolium* Braem, M.L.Baker & C.O.Baker (Lan and Albert 2011). Notably, there were two 5S rDNA sites on the same chromosome, such as *P. concolor* (Luo et al. 2017), *Brassica oleracea* L. (Ali et al. 2005), *O. cabulica* (Ali et al. 2005), *P. primulinum*, *P. liemianum*, *P. randsii*, *P. parishii*, *Paphiopedilum dianthum* T. Tang et F. T. Wang (Lan and Albert 2011), *Prunus cerasus* L. (Wang et al. 2022), and *Plantago maxima* Juss. ex Jacq. (Kovács et al. 2023).

Conversely, the 5S rDNA FISH signal position was diverse and abundant. The 5S rDNA signal has been found in the chromosome interstitial position from *Macroptilium bracteatum* (Nees & Mart.) Maréchal & Baudet (de Barros et al. 2023), *Deschampsia antarctica* E. Desv (Amosova et al. 2022), *Polemonium caeruleum* Linnaeus (Samatadze et al. 2023), in the chromosome distal position from *Pinus koraiensis* Siebold et Zuccarini (Cai et al. 2006), *Cannabis sativa* L. (Alexandrov et al. 2022), *Pseudotsuga menziesii* (Mlrb.) Franco (Amarasinghe and Carlson 1988), or in the chromosome proximal position from *Brassica rapa* L. (Campomayor et al. 2021), even as far away the chromosome from *Hedysarum setigerum* Turcz. (Yurkevich et al. 2022), and *P. cerasus* (Wang et al. 2022).

As a consequence, the 5S rDNA as a FISH probe was an excellent marker to label plant chromosomes in species and to distinguish closely related species among more than 300 plant species (listed in Supplementary Table 1), including more than 20 woody plant species (bold type in Supplementary Table 1). For example, in six species of Fabaceae: *Amorpha fruticose* L., *Styphnolobium japonicum* (L.) Schott, three species of *Robinia* L. (He et al. 2022a), *P. concolor* (Luo et al. 2017); in five *Pinus* L. species of Pinaceae (Cai et al. 2006); in four species of Oleaceae: *Fraxinus pennsylvanica* Marsh., two species of *Ligustrum* L., *Syringa oblata* Lindl. (Luo and Liu 2019); in two *Berberis* L. species of Berberidaceae (Liu and Luo 2019); in two species of Malvaceae: *Adansonia digitata* L. (Islam-Faridi et al. 2020), *Hibiscus mutabilis* L. (Luo and He 2021); and in Rutaceae of *Zanthoxylum armatum* DC. (Luo et al. 2018d, He et al. 2023).

However, the 5S rDNA as a FISH probe lacked discrimination (only stable two chromosomes with signal) in *Pistia stratiotes* L. of Araceae (Stepanenko et al. 2022); in *Chimonanthus campanulatus* R.H. Chang & C.S. Ding of Calycanthaceae (Luo and Chen 2019); in four *Citrullus* Schrad. species of Cucurbitaceae (Li et al. 2016); in cultural/wild *Hippophaë rhamnoides* ssp. *sinensis* and three *Hippophaë rhamnoides* L. cultivars of Elaeagnaceae (Luo et al. 2022a); in Fabaceae of six *Hedysarum* L. species (Yurkevich et al. 2022); in *Juglans regia* L. and *Juglans sigillata* Dode (Luo and Chen) of Juglandaceae; in four *Glechoma* L. species of Lamiaceae (Jang et al. 2016); in three *Bletilla* Rchb. f. species of Orchidaceae (Huan et al. 2022); in three *Zea* L. species of Poaceae (Han et al. 2003); in four *Citrus* L. species of Rutaceae (He et al. 2020), *Zanthoxylum bungeanum* Maxim. and *Z. armatum* (Hu et al. 2023); in four *Populus* L. species of Salicaceae (Xin et al. 2019); in five *Taxus* L. species of Taxaceae (He et al. 2022b).

Summarily, 5S rDNA has been confirmed to be a significant cytogenetic symbol by tandem formatting and giving in multicopy numbers with unusual chromosomal allocation. Moreover, these discoveries may afford a responsible method to survey the locations and number of rDNA diversifications among respective species and their relational accessions. Such findings also let us further understand the developmental and phylogenetic links of exampled species. The patent oligonucleotide FISH (Oligo-FISH) technology is widely problematic in related cytogenetic investigations compared to conventional FISH analysis because of its low cost (Beliveau et al. 2012). Moreover, based on available DNA sequencing data, oligonucleotide probes (Oligo-probe) have been developed and successfully applied to many plants (He et al. 2020, Xin et al. 2020, Luo et al. 2022a, b, He et al. 2023). Therefore, examining the 5S rDNA sites by Oligo-FISH would efficiently contribute to further cytogenetic research on the 64 plants. This study aimed to analyze and compare the polymorphism of the signal pattern of 5S rDNA among species and genera in the 64 plants examined, including 5S rDNA signal number, position, and intensity, and the chromosome number of each plant. Furthermore, several contributing factors caused by the diversity of the 5S rDNA signal pattern remain to be addressed.

## Materials and Methods

The species used for this experiment were chosen due to the discovery of karyotype realignments (Luo et al. 2017, Luo et al. 2018d, Liu and Luo 2019, Luo and Liu 2019, Luo ang Chen 2019, 2020, Luo and He 2021, Luo et al. 2022a, b, He et al. 2022a, b, c, 2023). Because these species were checked, their 5S rDNA has not yet been explored. Information on the seeds or seedlings of 64 plants (52 woody plants and 12 herbaceous plants belonging to 21 species, 18 genera, and 16 families) used in the present work is provided in Table 1. All 64 plants were collected from 23 counties or districts of six Chinese provinces.

**Table 1.**
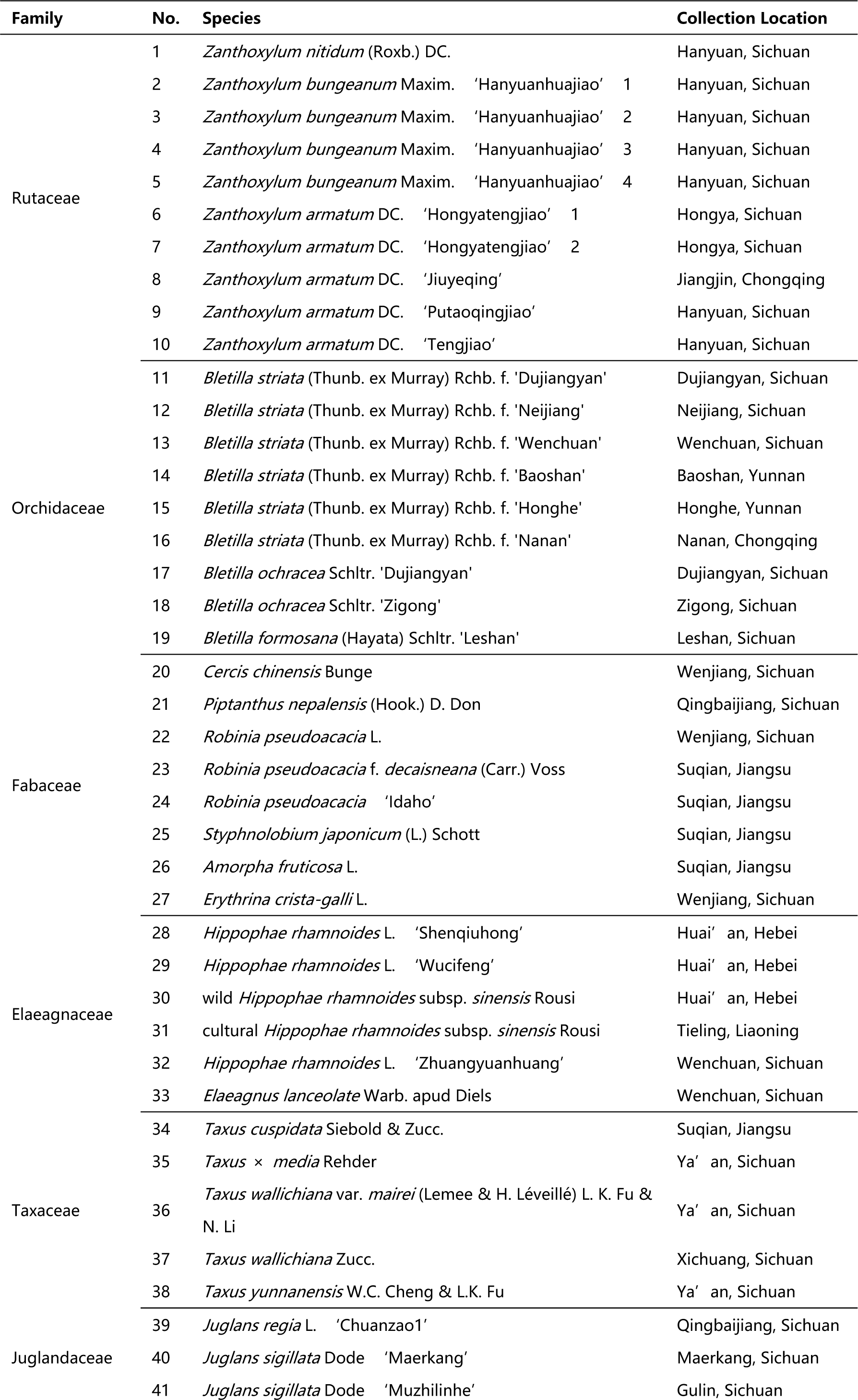
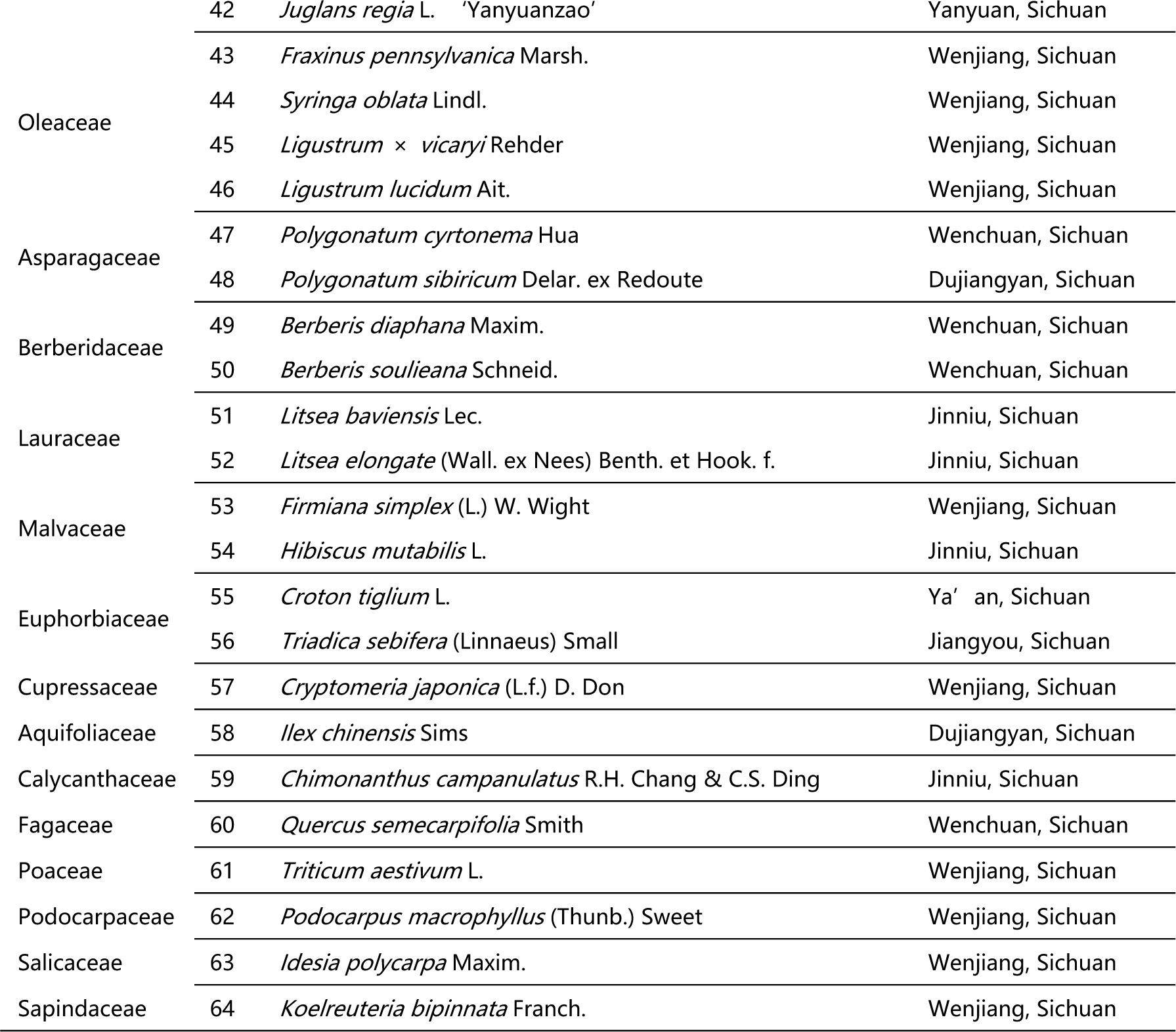
Details of all 64 plants used in this work.

### Chromosome and Oligonucleotide Probe Preparation

Converged seeds were germinated in Petri dishes with moistened filter paper and stayed at 25℃ in the daytime and at 18℃ in the night until the roots touched ∼2 cm in length. Next, the roots were cut. The collected seedlings were grown in soil at room temperature (15–25℃) until they had new roots and then cut again. The cut root tips were served with N2O (nitrous oxide) gas for 3–5 h, with processing time according to cell wall lignification and chromosome size. Afterward, the root tips were soaked in glacial acetic acid for 5 min and then in 75% ethyl alcohol. Chromosome preparation was executed based on the steps described by Luo et al. (2017). Since these procedures have been reported elsewhere, a brief description is available here. About 1 mm of the meristematic zone of the root tip (cut root cap) was in enzymolysis at 37℃ for 45 min by using pectinase and cellulase (the buffer 50 mL was covered 0.4324 g citric acid + 0.5707 g trisodium citrate, then 1 mL buffer + 0.02 g pectinase + 0.04 g cellulase). The enzymes were yielded by Kyowa Chemical Products Co., Ltd. (Osaka, Japan) and Yakult Pharmaceutical Ind. Co., Ltd. (Tokyo, Japan). The enzymes were then blended into suspension for dropping onto clean slides. These slides were air dried at room temperature and examined using an Olympus CX23 microscope (Olympus, Tokyo, Japan).

The oligoprobe of 5′ TCAGAACTCCGAAGTTAAGCGTGCTTGGGCGAGAGTAGTAC 3′ (41 bp) was initially reported in *P. nepalensis* (*P. concolor*, former name in Flora of China) of Fabaceae (Luo et al. 2017). Then, it has been steadily executed in two *Berberis* species of Berberidaceae (Liu and Luo 2019), *C. campanulatus* of Calycanthaceae (Luo and Chen 2019), and in cultural/wild *H. rhamnoides* ssp. *sinensis* and three *H. rhamnoides* cultivars of Elaeagnaceae (Luo et al. 2022a), in *A. fruticose* and *S. japonicum* of Fabaceae and three *Robinia* species (He et al. 2022a), two *Juglans* species of Juglandaceae (Luo and Chen 2020), *A. digitata* and *H. mutabilis* of Malvaceae (Islam-Faridi et al. 2020, Luo and He 2021), *F. pennsylvanica* of Oleaceae, two species of *Ligustrum* and *S. oblata* (Luo and Liu 2019), three *Bletilla* species of Orchidaceae (Huan et al. 2023), two *Zanthoxylum* species of Rutaceae (Luo et al. 2018, He et al. 2023), and five *Taxus* species of Taxaceae (He et al. 2022b). This oligoprobe was produced by Sangon Biotech Co., Ltd. (Shanghai, China) and conducted simultaneously in a single round of FISH. The oligo-probe was 50-labeled with 6-carboxy-fluorescein (6-FAM; absorption /emission wavelengths 494 nm/518 nm; green).

### FISH Hybridization

Slides with well-spread chromosomes were used to hybridize. Chromosome samples first experienced a series of reinforcement (4% paraformaldehyde, room temperature, 10 min), dehydration (75%, 95%, and 100% ethanol at room temperature for 5 min), denaturation (deionized formamide at 80℃ for 2 min), and once again dehydration (75%, 95%, and 100% ethanol, at −20℃ for 5 min), and then hybridization (0.375 μL of 5S rDNA, 4.675 μL of 2∈ SSC, and 4.95 μL of ddH_2_O in total 10 μL hybridization mixture) for 2 h in an incubator at 37℃. Then, hybridized spreads were rinsed with 2∈ SSC and ddH_2_O twice for 5 min at room temperature and air-dried. Finally, the spreads were counterstained with 4,6-diamidino-2-phenylindole (DAPI, Vector Laboratories, Inc., Burlingame, CA, USA) for 5 min, according to the step described by Luo et al. (2017). Chromosomes were traced using an Olympus BX-63 microscope (Olympus Corporation, Tokyo, Japan), and FISH photographs were taken using a DP-70 CCD camera allocated to the microscope.

### Karyotype Analysis

Raw data were executed using the Photoshop CC 2015 (Adobe Systems Inc., San Jose, CA, USA) and DP Manager (Olympus Corporation, Tokyo, Japan) software. More than eight slides of each plant were observed, and at least 15 well-spread cells were executed to count the chromosome number and length. All chromosomes examined were assembled from longest to shortest. The chromosome ratio was controlled by the length of the longest chromosome to that of the shortest chromosome. Exhaustive and deep karyotype analysis could not be conducted because of the obscure centromere position and small chromosome size of many of the species.

## Results

### Karyotype Analysis Revealed Differences among 64 Plants

We performed FISH analysis to visualize the chromosomal distribution of 5S rDNA in 64 plants, as shown in Figure 1 (A1–A16), Figure 2 (B1–B16), Figure 3 (C1–C16), and Figure 4 (D1–D16). We cut each chromosome distribution of the 5S rDNA in Figures 1–4 and aligned them for display, as shown in Figure 5 (A1–A16), Figure 6 (B1–B16), Figure 7 (C1– C16), and Figure 8 (D1–D16). For 19 species from 13 families, this is the first time that 5S rDNA testing has been analyzed (A1, A8–A11, A13–A16, B11, B14, C12, C14–C16, D6–D12, and D14–D15).

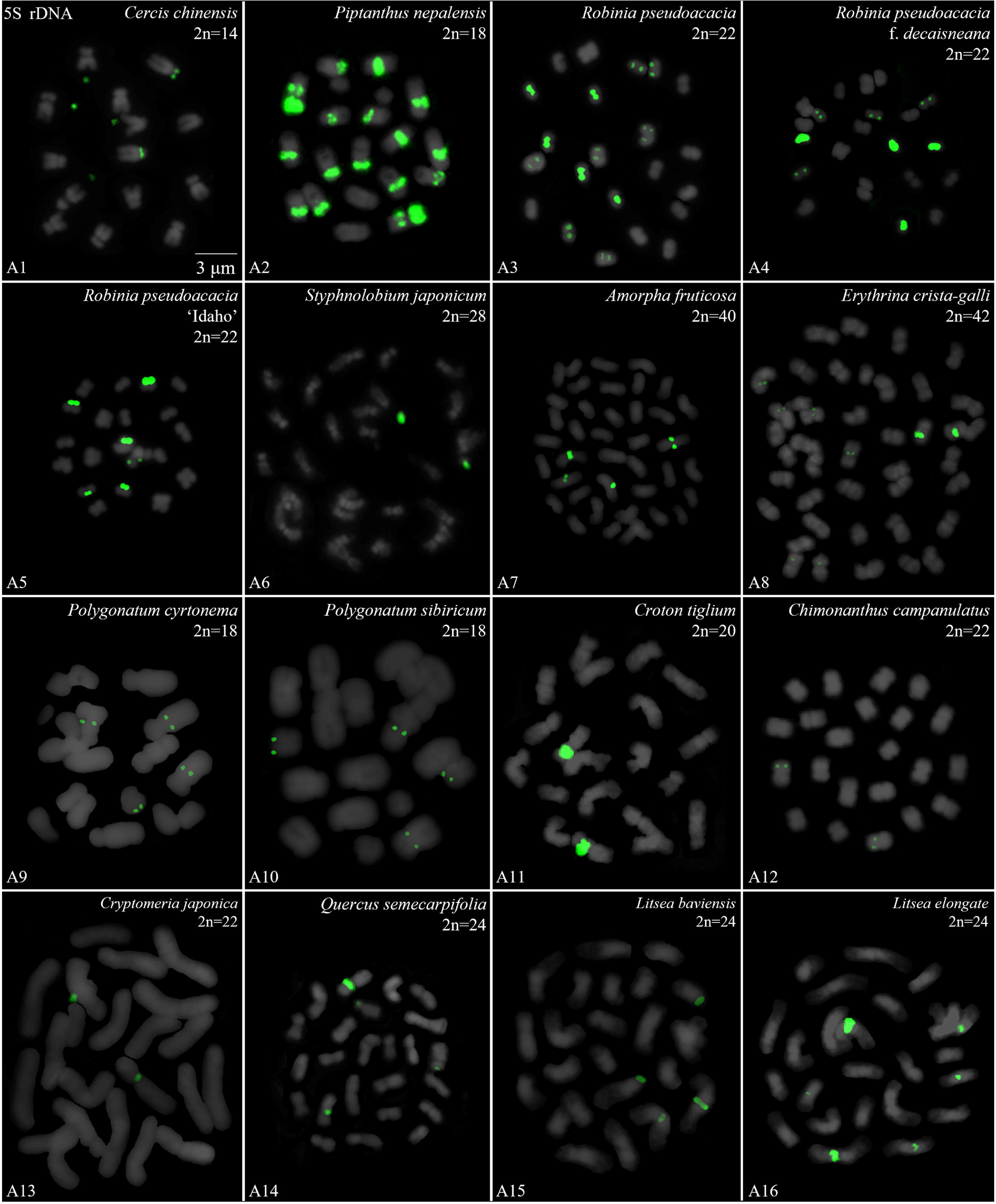

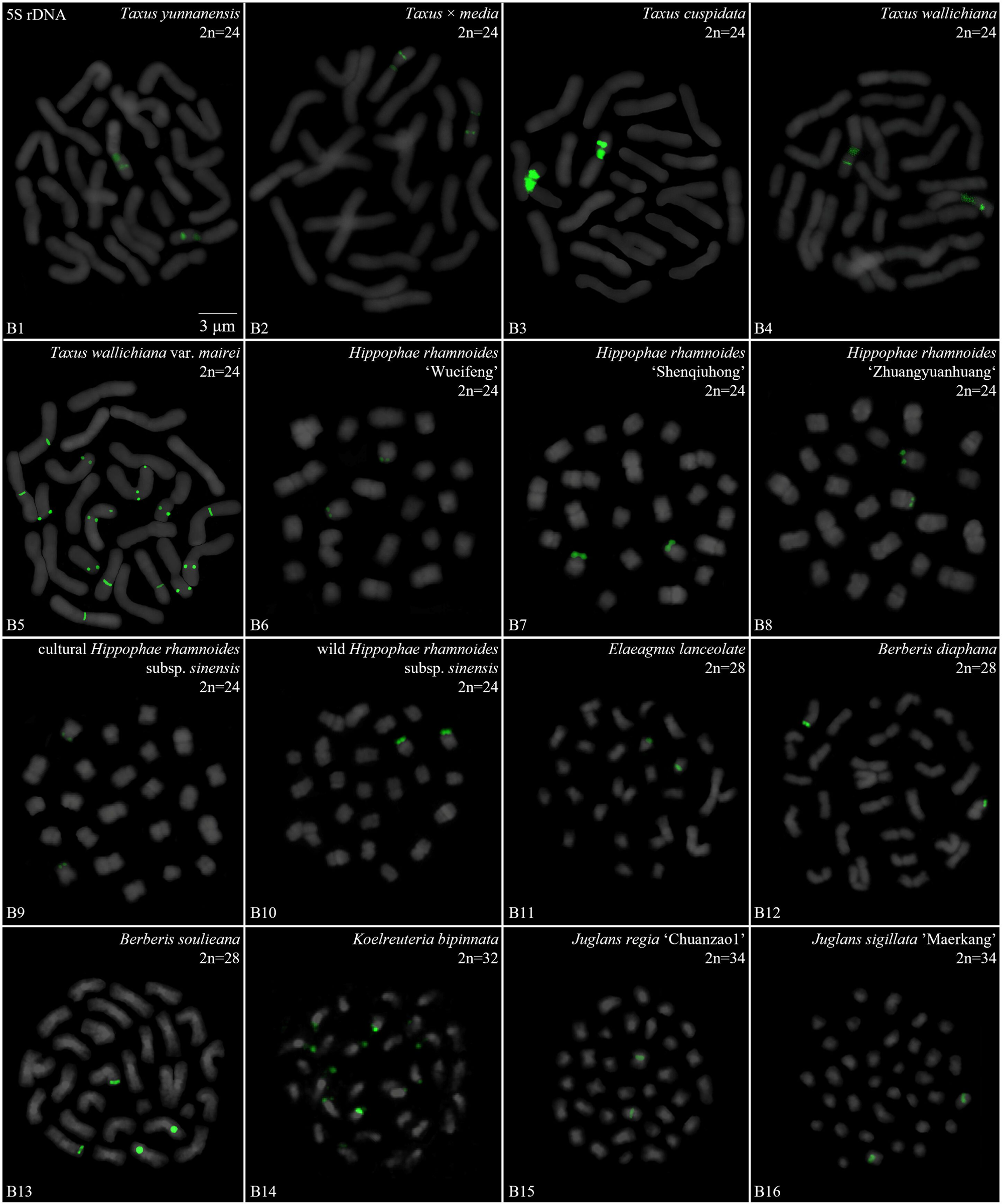

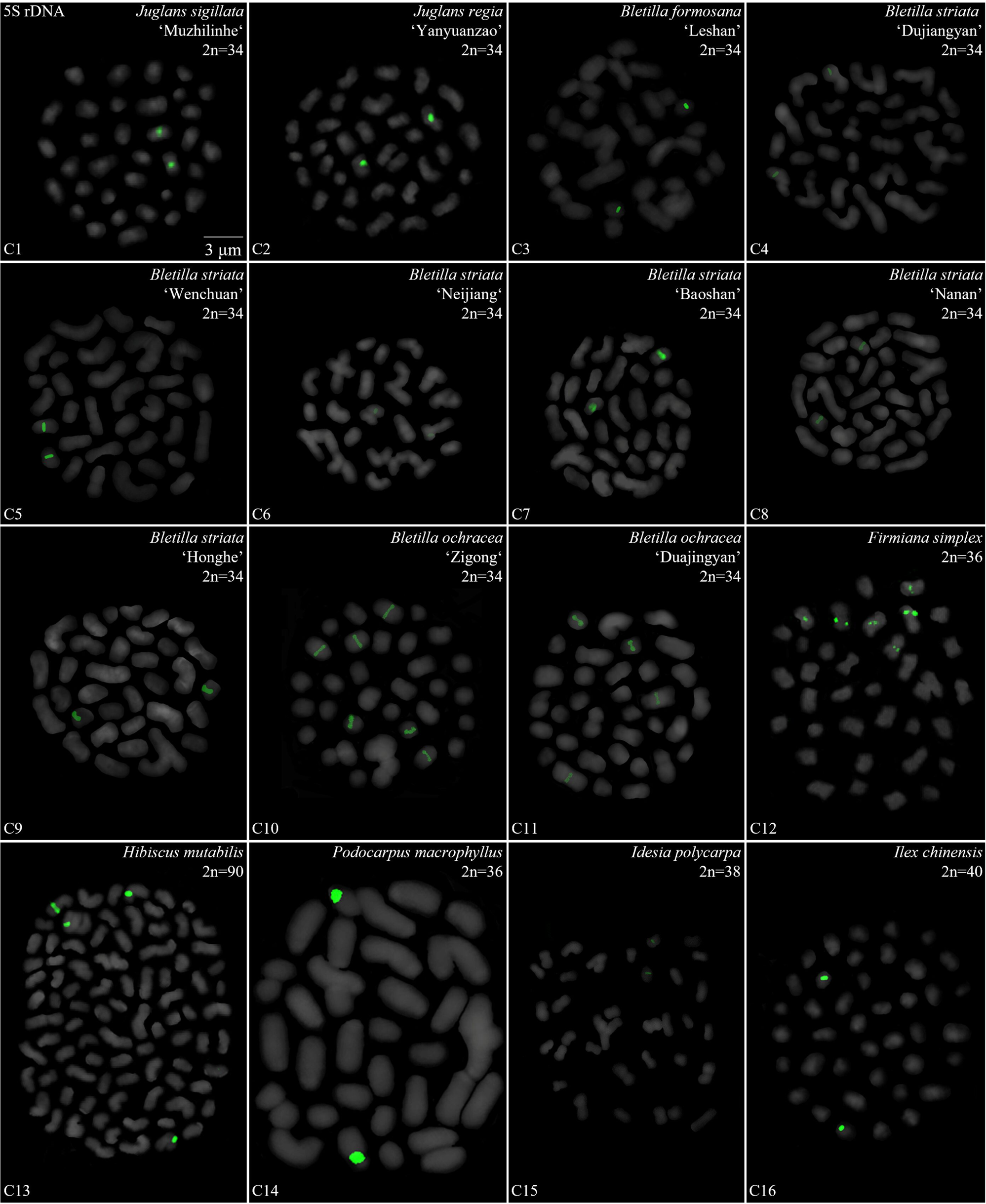

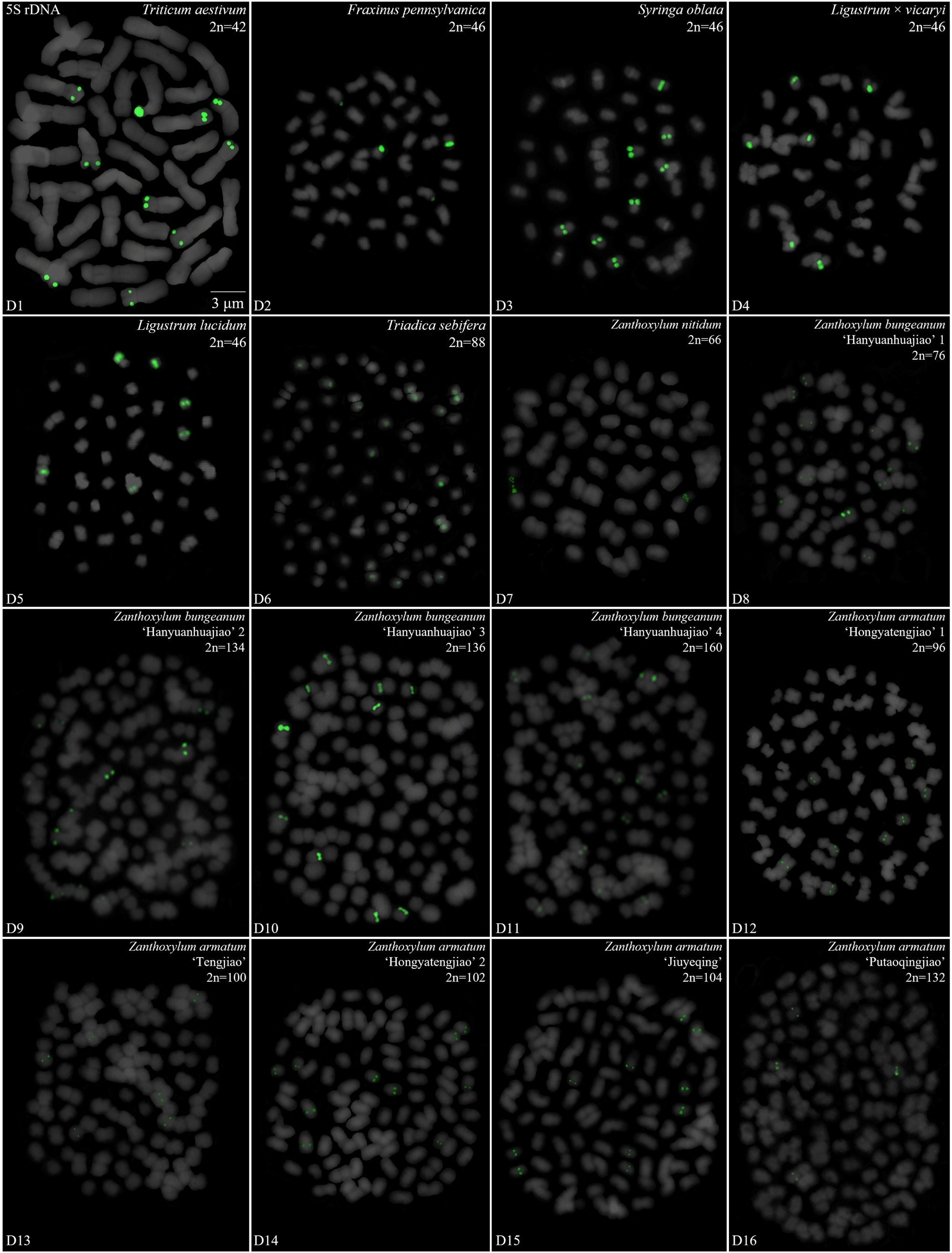

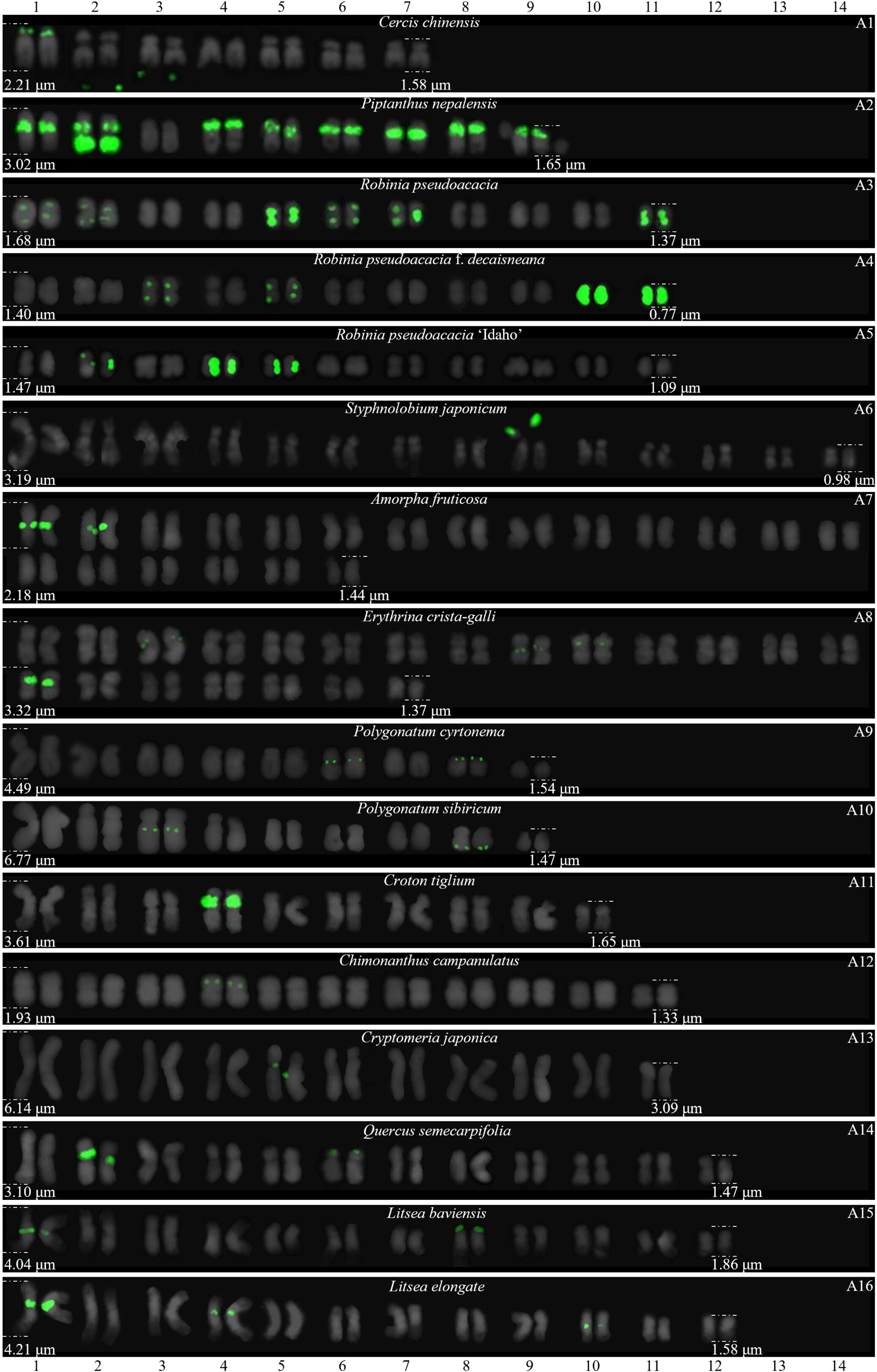

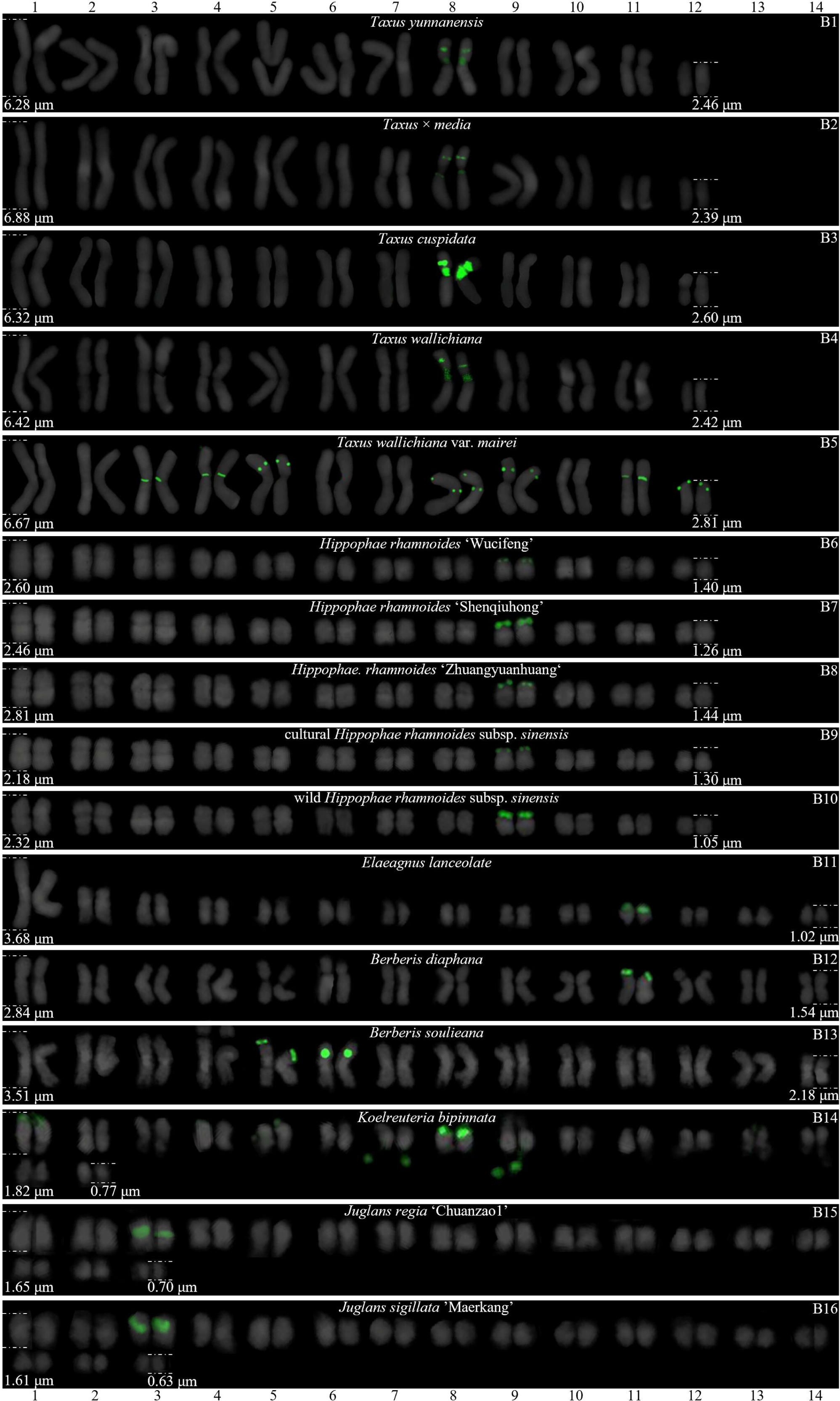

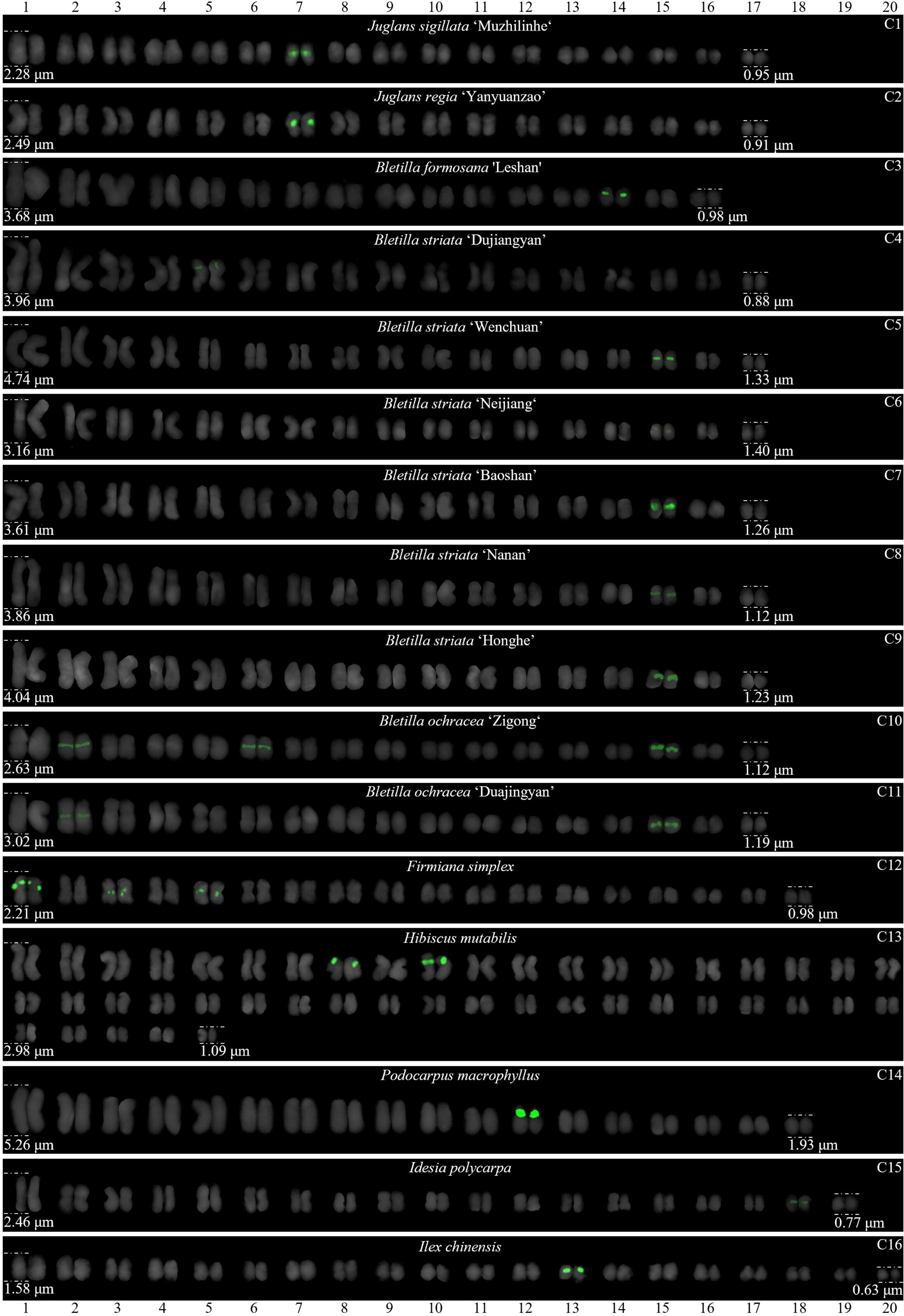

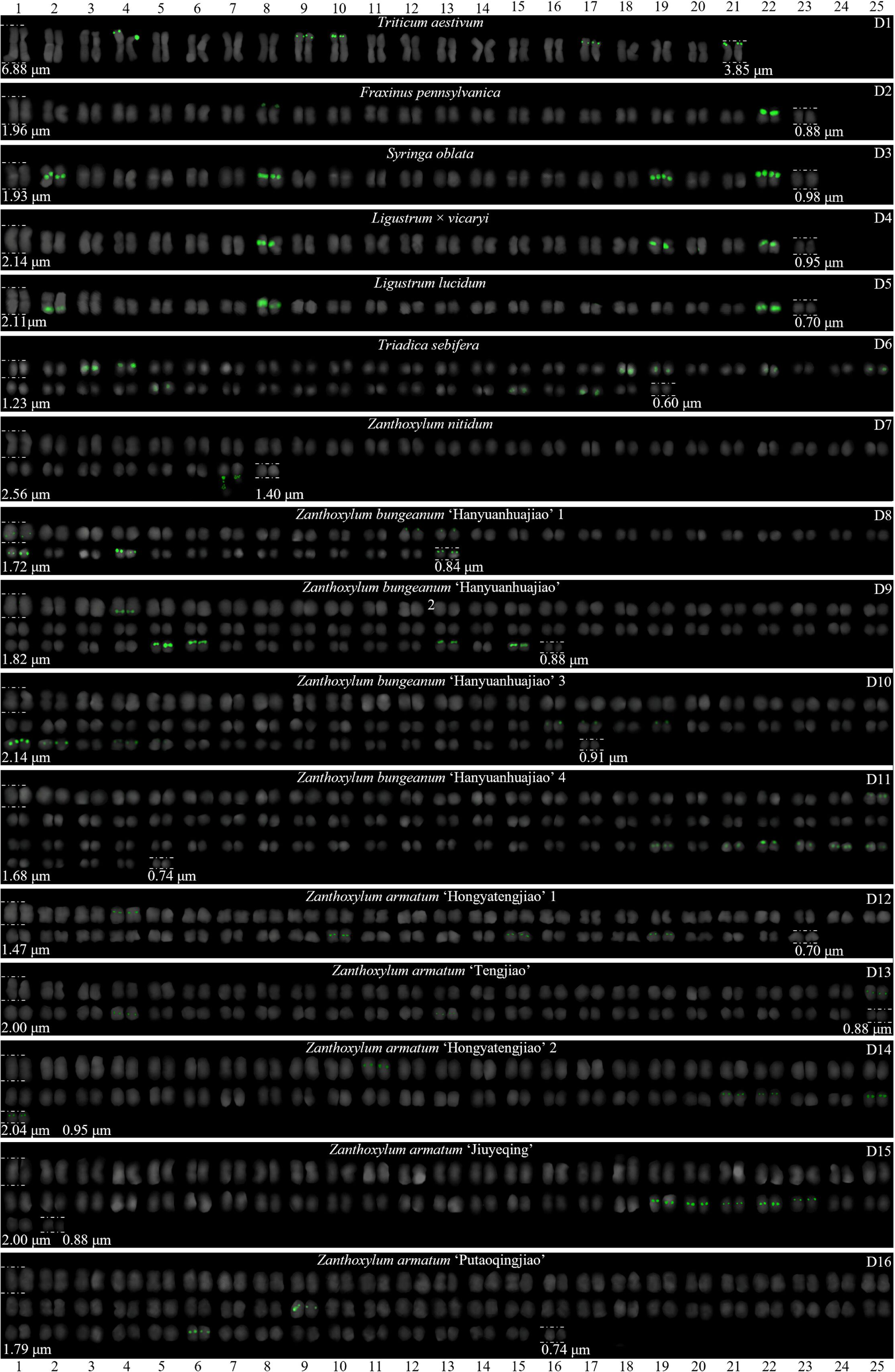

The chromosome numbers and lengths for the considered species are sorted in Table 2. The chromosome numbers in the 64 plants ranged from 14 (*C. chinensis*, A1) to 160 (*Z. bungeanum* ‘Hanyuanhuajiao’ 3, D11). Thirteen plants possessed 24 chromosomes (one-fifth), whereas another 13 plants possessed 34 chromosomes (one-fifth). The chromosome number of the three species was analyzed for the first time here (bold type in Table 2): *P. sibiricum* (2n = 18), *I. chinensis* (2n = 40), and *T. sebifera* (2n = 88). The longest chromosome length of each plant ranged from 1.23 μm (*T. sebifera*, D6) to 6.88 μm (*T.* × *media*, B2; *T. aestivum*, D1). The shortest chromosome length of each plant ranged from 0.63 μm (*J. sigillata* ‘Maerkang,’ B16) to 3.85 μm (*T. aestivum*, D1). Thirty-seven plants (58%) had a chromosome length of less than 3 μm, thus falling into the small chromosome category. Detailed and deep karyotype analysis was not conducted because of the unclear position of centromeres and the small size of chromosomes in many of the exampled plants, such as long/short arm length and karyotype formula.

**Table 2.**
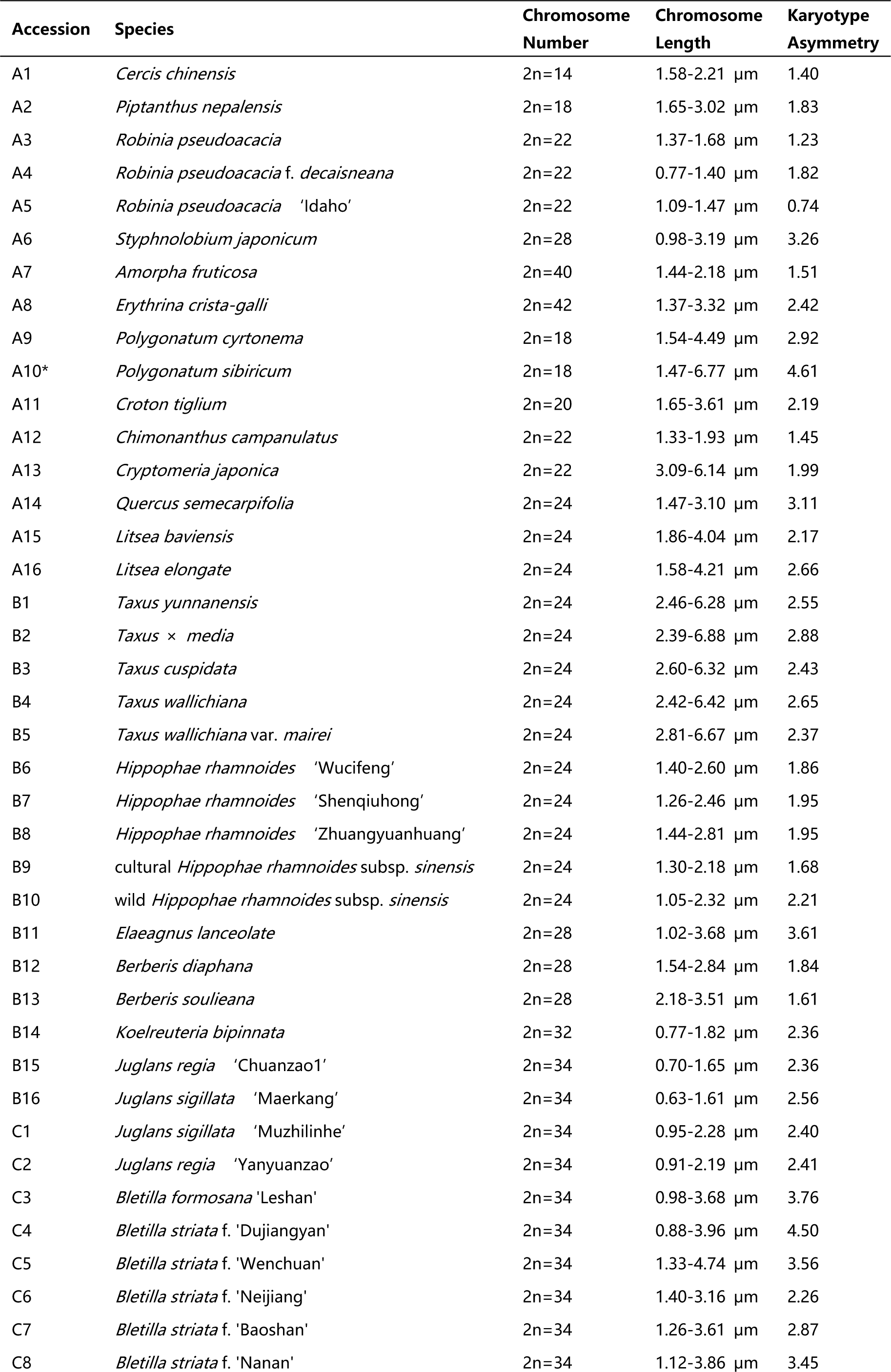
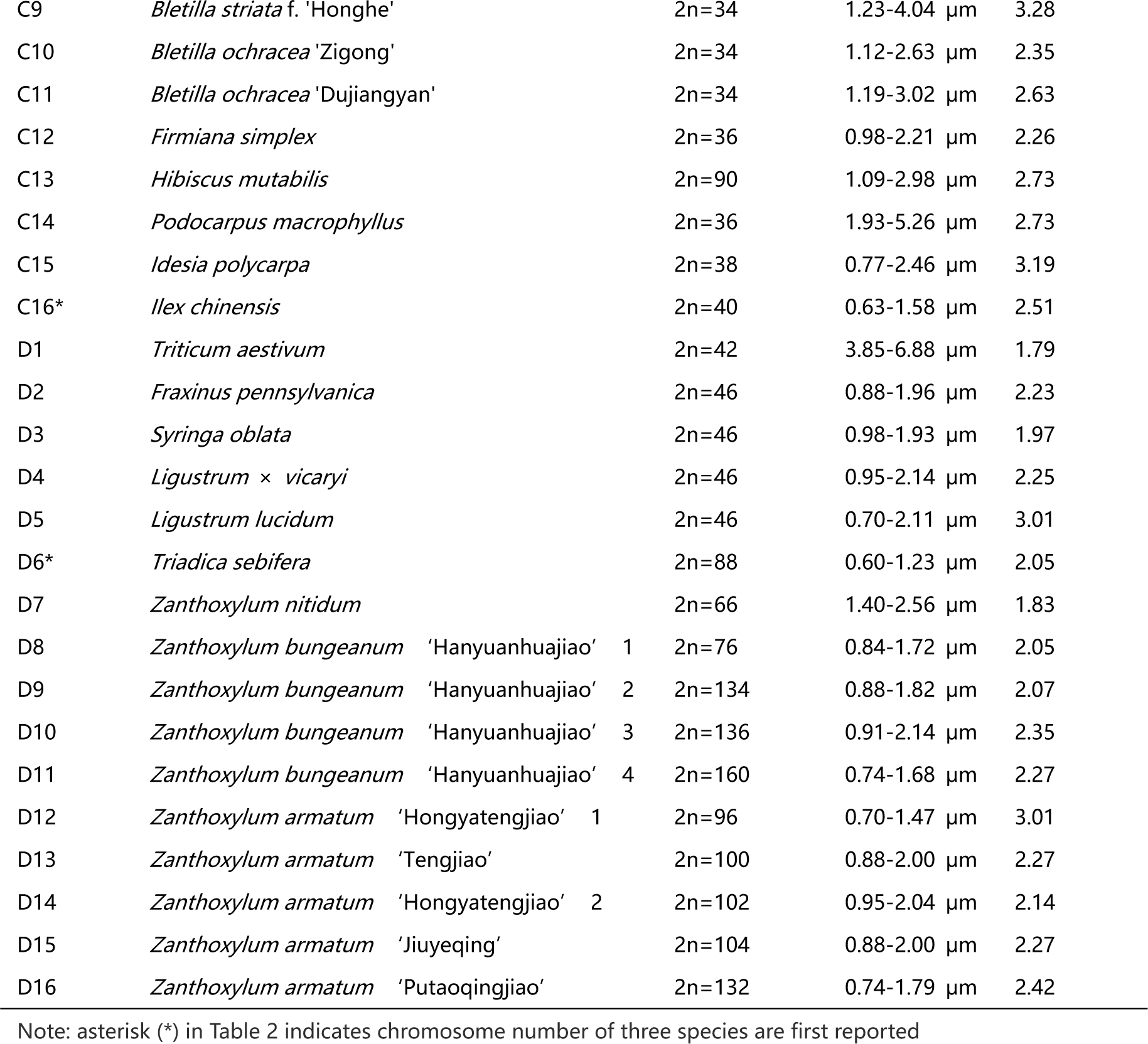
Chromosome number and length of the 64 plants used in this work.

Karyotype asymmetry (Table 2) was assessed using the longest to shortest chromosome length ratio. The largest ratio was 4.61 in *P. sibiricum* (A10), while the smallest ratio was 0.74 in *R. pseudoacacia* ‘Idaho’ (A5). The ratio for 35 plants ranged from 2 to 3 (55%), while that of 16 plants ranged from 1 to 2 (25%), and 10 plants ranged from 3 to 4 (16%). The ratio was greater than 4 for two plants: *P. sibiricum* (A10) and *B. striata* f. ‘Dujiangyan’ (C4), while the ratio was less than 1 for *R. pseudoacacia* ‘Idaho’ (A5). These results indicate that abundant differences exist among 44 of the considered species.

### The Diverse Signal Patterns of 5S rDNA Reveal the Complex Genome Architecture

Different types of ideograms for the 64 plants were drawn based on the FISH karyograms shown in Figures 5–8 to better investigate the diversity of 5S rDNA (Figures 9–12). The first diversity is signal location. Proximal signals were observed at several chromosomes in 39 plants (61%), while distal signals were observed at several chromosome terminus in 30 plants (47%). Interstitial signals were observed at several chromosomes in 21 plants (33%), while distal signals deviated from the chromosome in 4 plants (6%, the fourth class): *C. chinensis* (A1), *S. japonicum* (A6), *K. paniculata* (B14), and *Z. nitidum* (D7). The second diversity is the signal number. The largest number of chromosomes with a 5S rDNA signal was 18 in *T. sebifera* (D6), while the smallest number was 2 in 31 plants (48%). The number of chromosomes with a 5S rDNA signal for 11 plants ranged from 10 to 16 (17%), while that of 21 plants ranged from 4 to 8 (33%). The ratio of chromosomes with a 5S rDNA signal to total chromosome was assessed to signal cover. The largest ratio was 0.89 in *P. nepalensis* (A2), while the smallest ratio was 0.03 in *Z. nitidum* (D7) and *Z. armatum* ‘Putaoqingjiao’ (D16). The ratio for 50 plants ranged from 0.03 to 0.20 (78%), while the ratio for nine plants ranged from 0.20 to 0.50 (14%). The ratio for only two plants ranged from 0.50 to 0.89 in *R. pseudoacacia* (A3) and *T. wallichiana* var. *mairei* (B5).

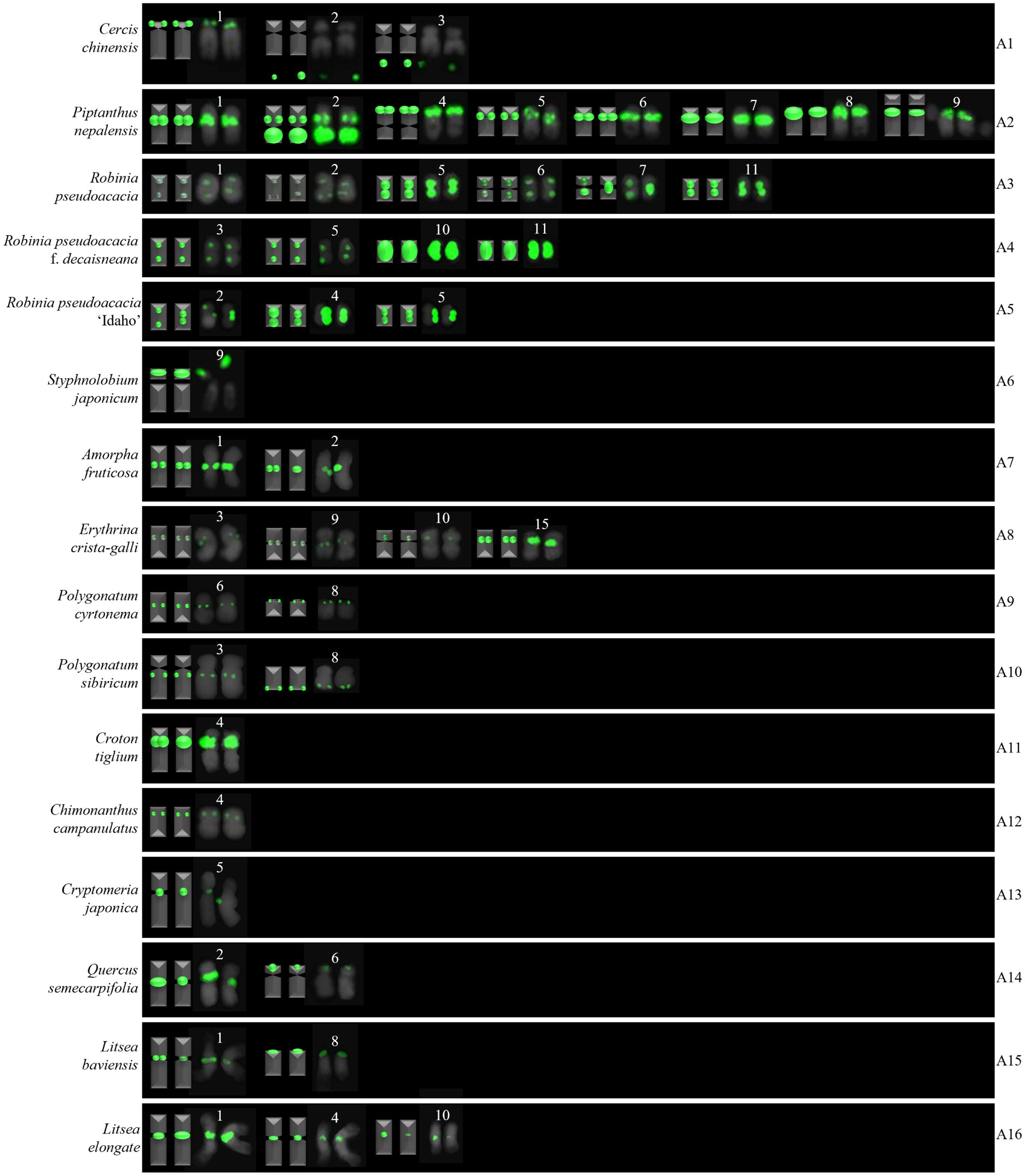

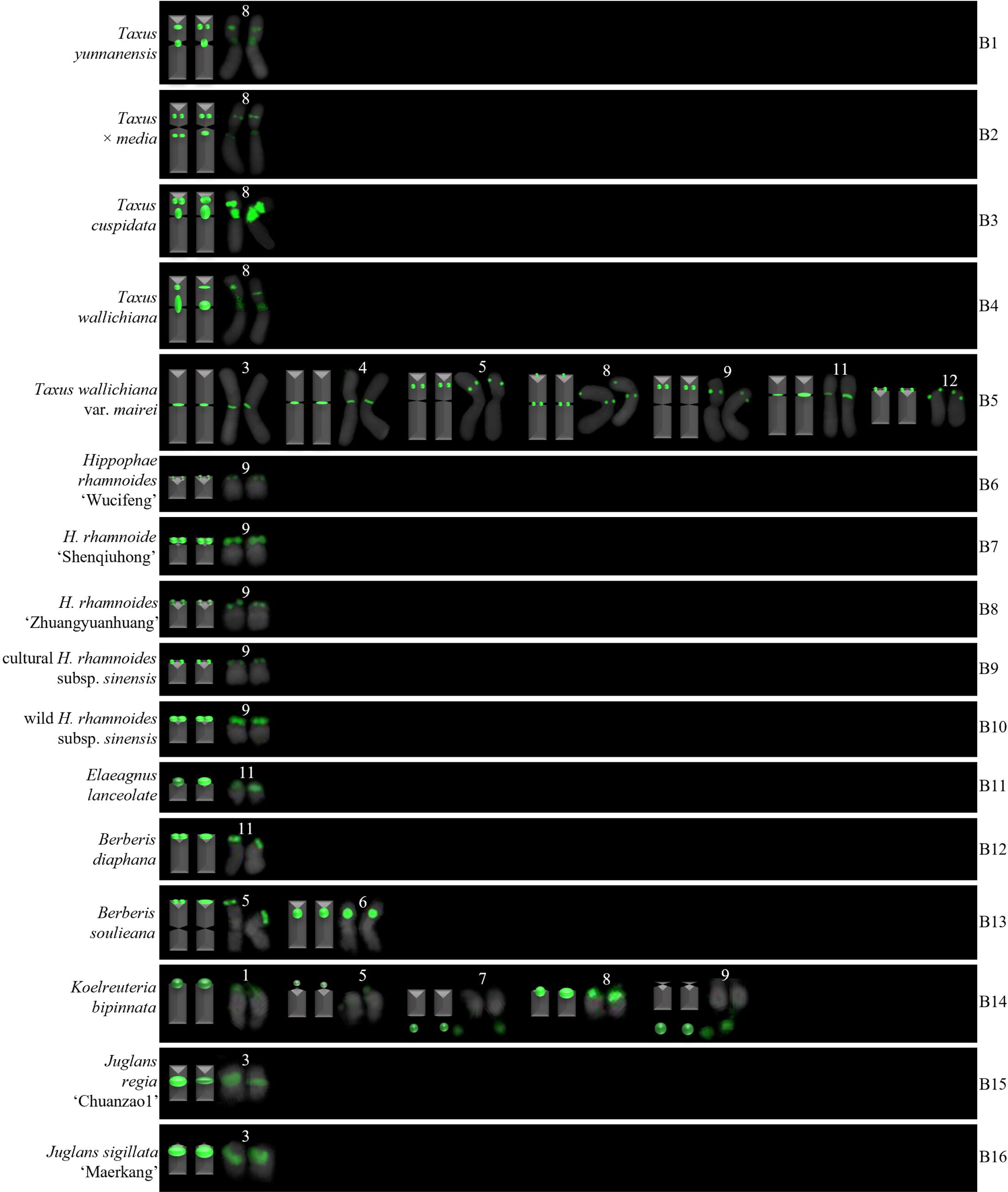

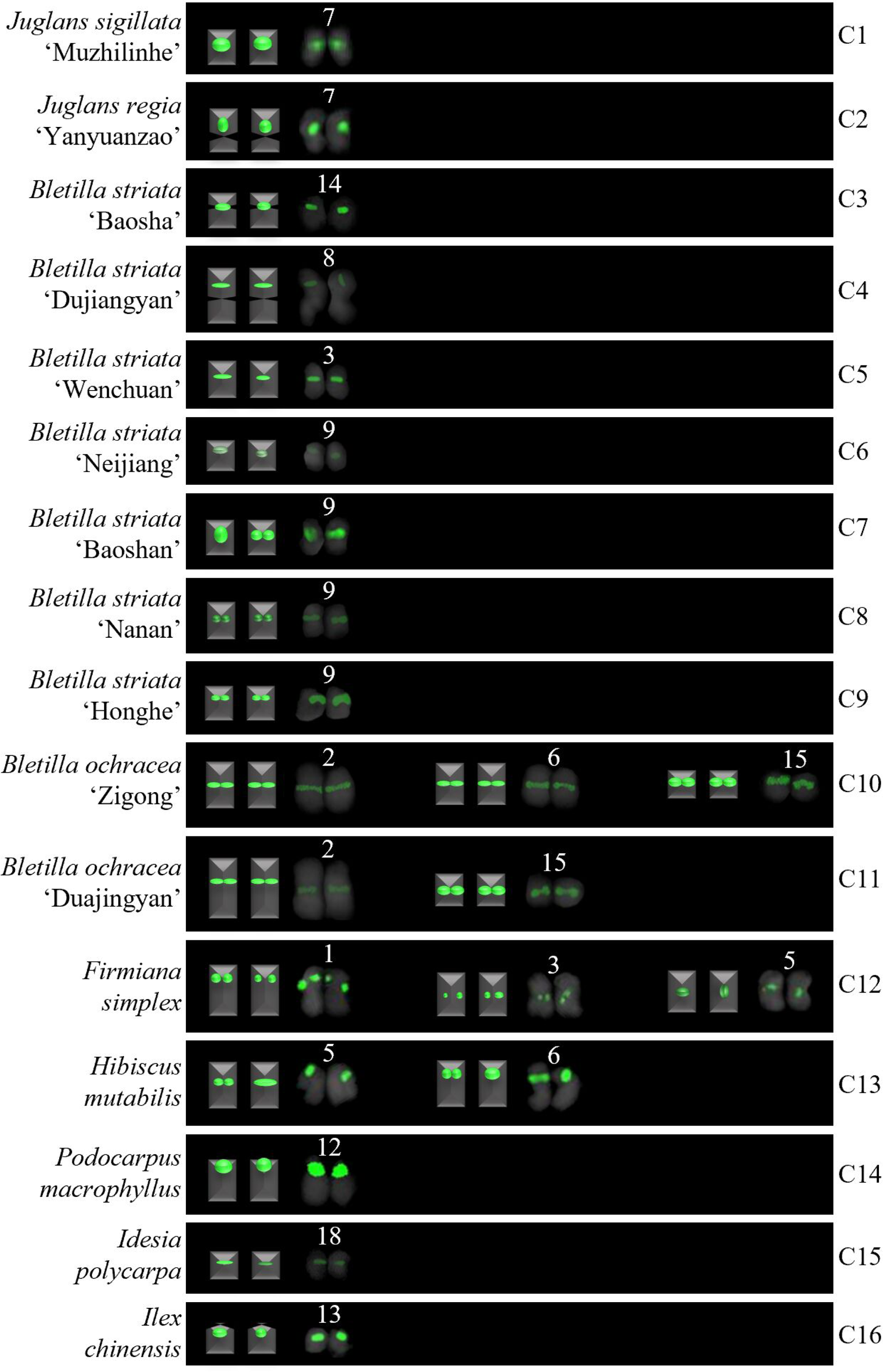

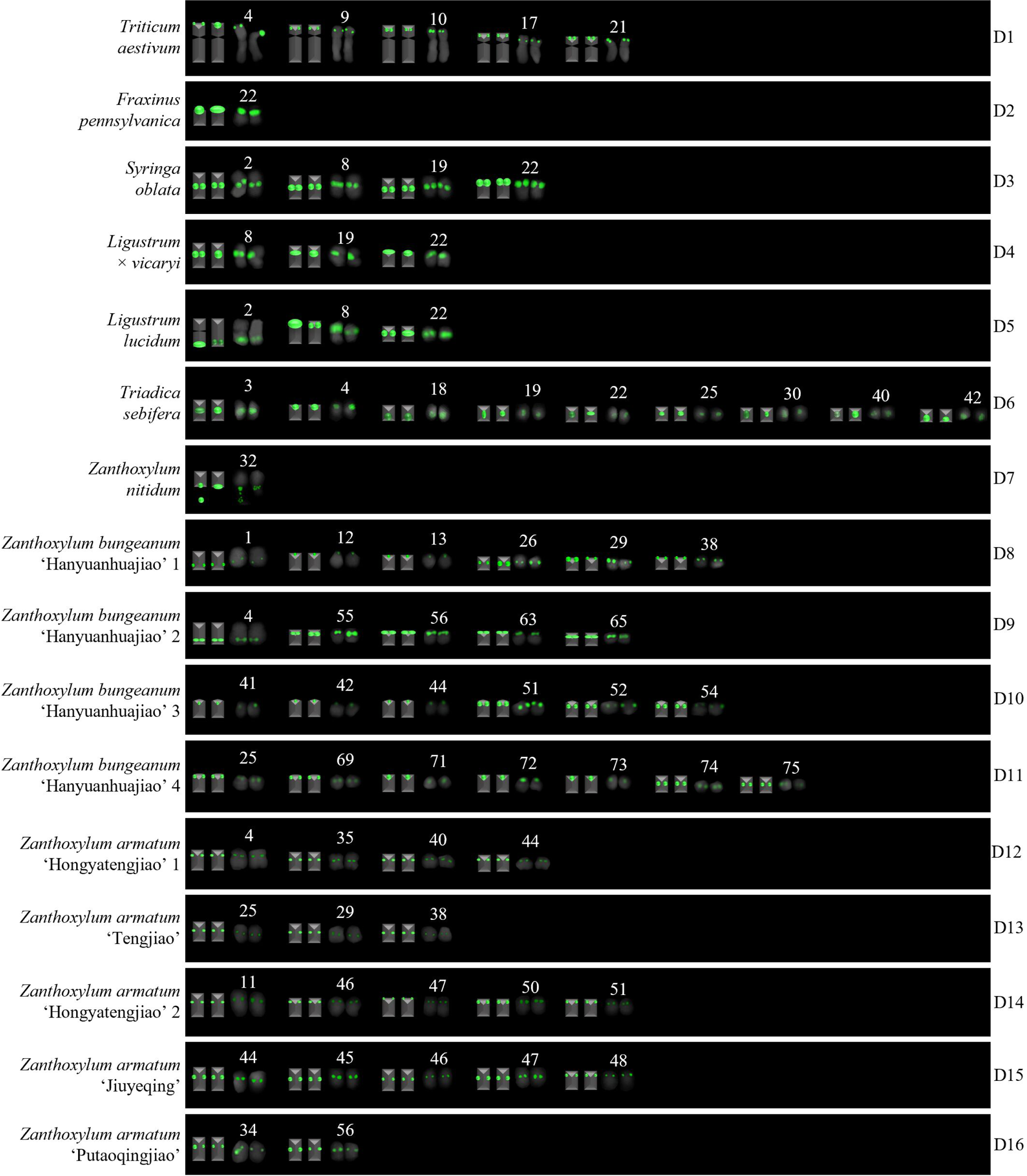

Furthermore, we summarized the results in Figures 9–12 to produce the 5S rDNA signal pattern in Figure 13. The results for 64 plants belonging to 20 families are shown, including 10 plants in Rutaceae (gray), nine in Orchidaceae (light blue), eight in Fabaceae (cyan), six in Elaeagnaceae (light yellow), five in Taxaceae (light pink), four in Juglandaceae (green), four in Oleaceae (orange), two in Asparagaceae (yellow), two in Berberidaceae (blue), two in Euphorbiaceae (red), two in Malvaceae (magenta), two in Lauraceae (pink), and one in each of Aquifoliaceae, Calycanthaceae, Cupressaceae, Podocarpaceae, Fagaceae, Poaceae, Salicaceae, and Sapindaceae, respectively. Except for Aquifoliaceae, Asparagaceae, Calycanthaceae, Cupressaceae, Elaeagnaceae, Juglandaceae, Orchidaceae, Podocarpaceae, and Salicaceae, which each presented a single signal pattern type, the other families (Berberidaceae, Euphorbiaceae, Fagaceae, Fabaceae, Lauraceae, Malvaceae, Oleaceae, Poaceae, Rutaceae, Sapindaceae, and Taxaceae) all presented at least two sites.

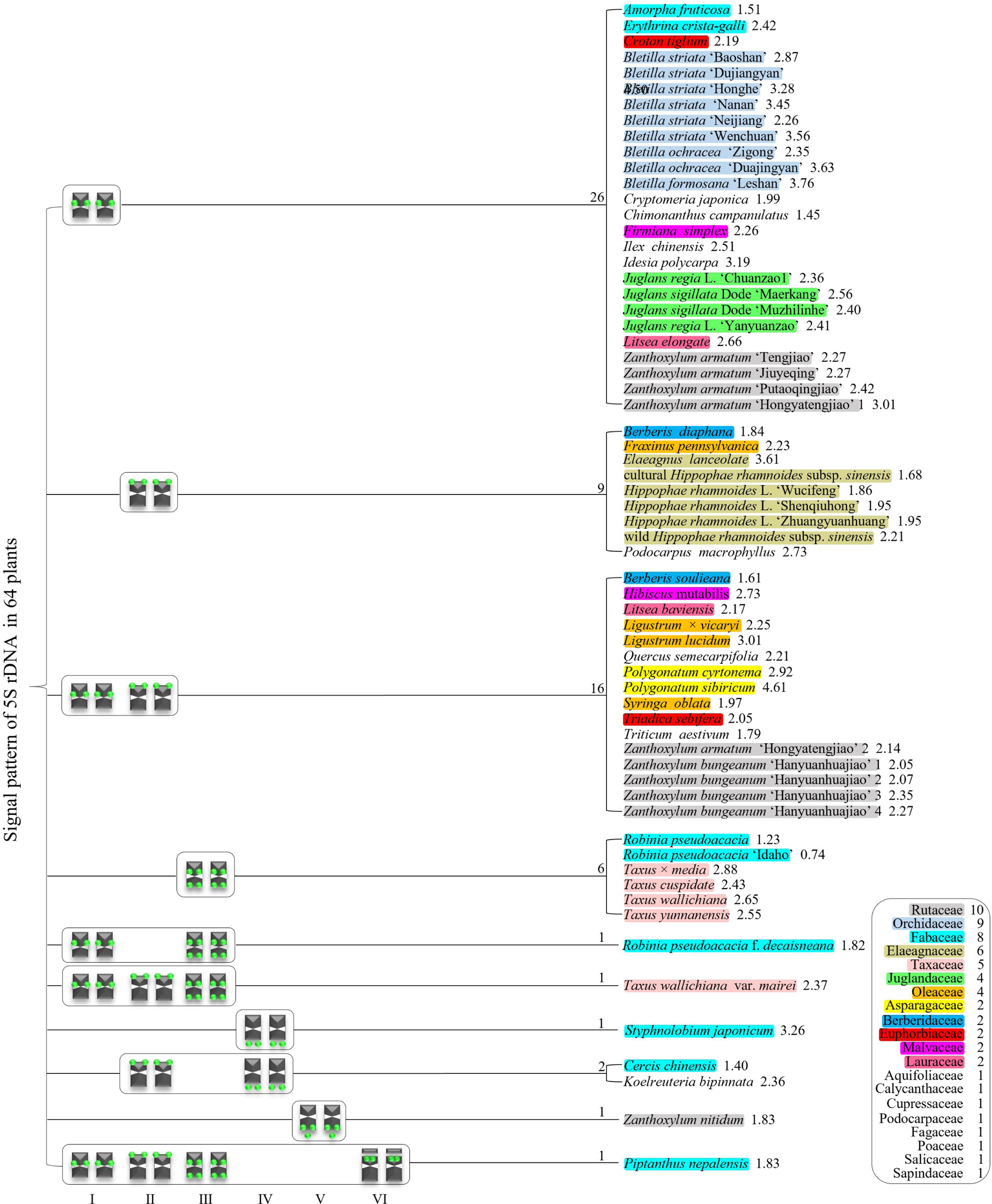

As shown in Figure 13, there were six 5S rDNA signal pattern types in total: type I, chromosome includes the proximal signal location; Type II, chromosome includes the distal signal location; Type III, chromosome includes the proximal and distal signal locations; Type IV, chromosome only includes signal outside of the chromosome; Type V, chromosome includes the distal signal location and signal outside of the chromosome; and Type VI, satellite chromosome includes distal signal location. These types of signal patterns indicate that there is abundant diversity in the 5S rDNA signal arrangement.

All 64 plants had the 10 signal pattern types or type combinations shown in Figure 13. The 26 plants only possessed signal pattern type I; nine plants only possessed signal pattern type II; 16 plants had a combination of type I + type II; six plants only had type III; *R. pseudoacacia* f. *decaisneana* had a combination of type I + type III; *T. wallichiana* var. *mairei* had a combination of type I + type II + type III; *S. japonicum* only had type IV; *C. chinensis* and *K. bipinnata* had a combination of type II + type IV; *Z. nitidum* only had type V; and *P. nepalensis* had a combination of type I + type II + type III + type VI.

There were diverse signal patterns of 5S rDNA among 44 species, indicating a complex genome architecture. For example, considering (*i*) in Rutaceae, four varieties of *Zanthoxylum* had type I, but five varieties of *Zanthoxylum* had a combination of type I + type II, and *Z. nitidum* had type V. (*ii*) In Fabaceae, A. fruticose and *E. crista-galli* had type I, but *R. pseudoacacia* and *R. pseudoacacia* ‘Idaho’ had type III, *R. pseudoacacia* f. *decaisneana* had combination of type I + type III, *S. japonicum* only had type IV, *C. chinensis* had a combination of type II + type IV, and *P. nepalensis* had a combination of type I + type II + type III + type VI. (*iii*) In Taxaceae, four species of *Taxus* had type III, but *T. wallichiana* var. *mairei* had a combination of type I + type II + type III. (*iv*) In Oleaceae, *F. pennsylvanica* had type II, but *L. lucidum*, *L.* × *vicaryi* and *S. oblata* had a combination of type I + type III. (*v*) In Berberidaceae, *B. diaphana* had type II, but *B. soulieana* had a combination of type I + type II. (*vi*) In Malvaceae, *F. simplex* had type I, but *H. mutabilis* had a combination of type I + type II. (*vii*) In Lauraceae, *L. baviensis* had type I, but *L. baviensis* had a combination of type I + type II.

### Proposed Origin of 5S rDNA Signal Diversity

Based on Figures 9–13, the proposed origin of the 5S rDNA signal diversity is illustrated in Figure 14. There are three major groups. (*i*) Signal number: signal number increases probably caused by the insertion of transposable elements, inversion, or translocation, and transposition events. Signal number decrease probably by elimination from the genome or conversion into pseudogenes and then loss, fusion event, and polyploidization followed by DNA sequence loss. (*ii*) Signal location on distal chromosome: polyploidization-related tendency toward the terminal location from an interstitial location. Signal location on proximal chromosome: interstitial or centromeric, a massive trend apparent in species with a single locus. The end signal deviation from the chromosome is probably caused by chromosome satellite, while transposon-mediated transpositional events and gene silencing probably cause end signal loss. (*iii*) Signal size: Signal size increase is probably caused by artificial selection, environmental induction, unequal crossing over recombination, illegitimate recombination, and duplication. The normal signal size indicates chromosome conservation. The signal size decrease is probably caused by self-incompatibility non-allelic homologous recombination. In brief, the signal number, location, and size variations were probably caused by chromosome rearrangement (deletion, duplication, inversion, and translocation), polyploidization, self-incompatibility, and chromosome satellites.

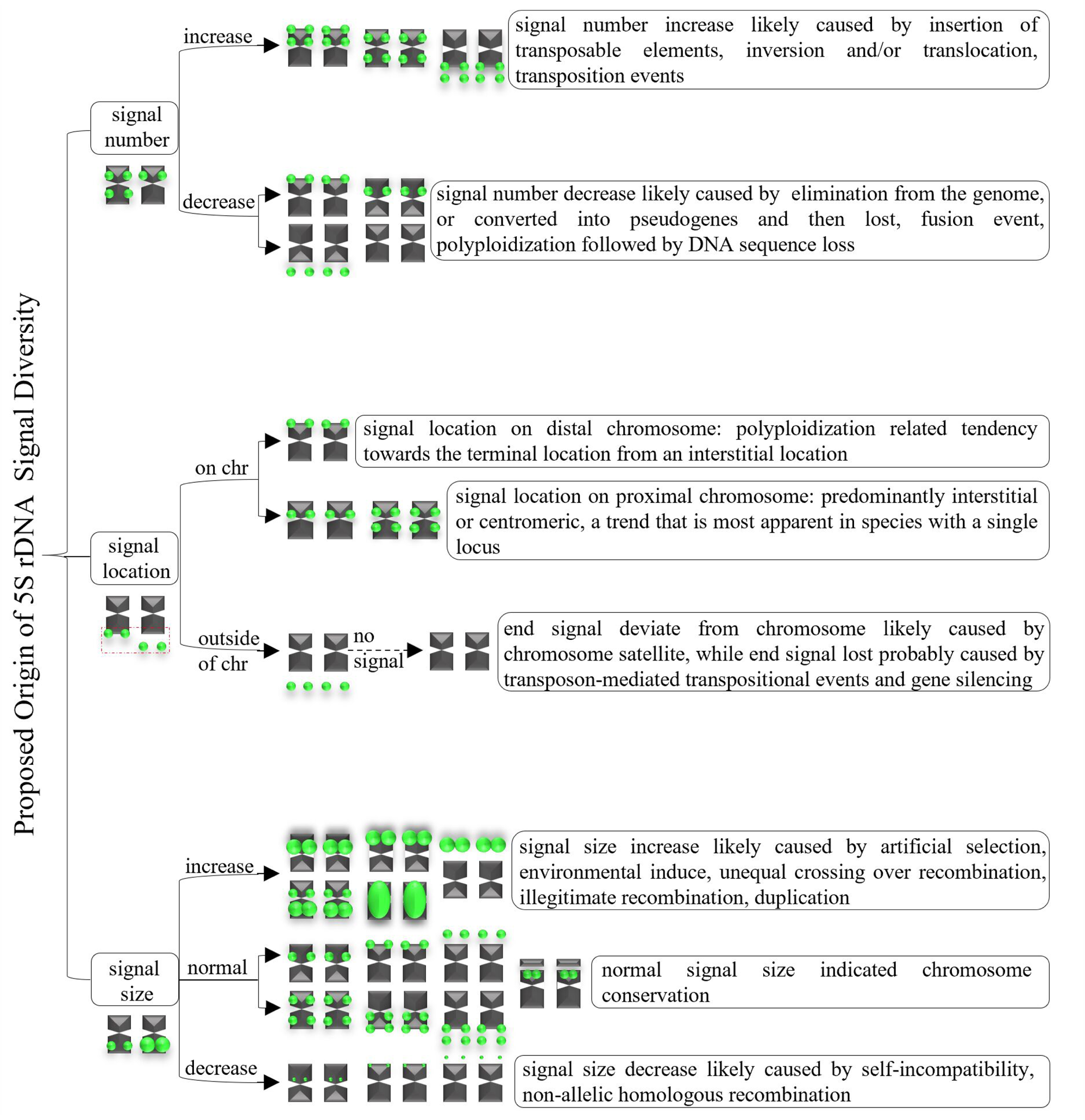

## Discussion

### Karyotype Analysis of the 64 Plants

Traditional karyotype analysis involves counting chromosome numbers, determining centromere location, and measuring chromosome length and long/short arm ratio. Based on this, the karyotype formula and cytotype are obtained to compare whether the species is evolving (Alberts et al. 2002). In this study, the chromosome numbers in the 64 plants ranged from 2n = 14 to 160. Most chromosome numbers are consistent with previous studies (Sun 1996, Chen and Zhou 2005, Luo et al. 2017, Sergeeva et al. 2017, Liu and Luo 2019, Luo and Liu 2019, Wang et al. 2019, He et al. 2022a, b, c, Yang et al. 2023). Only nine species had different chromosome numbers. In addition, the chromosome number of three species was the first time to be analyzed here: *P. sibiricum* (2n = 18), *I. chinensis* (2n = 40), and *T. sebifera* (2n = 88) in this study. More related information on *T. sebifera* was found in the chromosome number of the previous research. A stable chromosome number was found in the genus *Ilex*: *Ilex crenata* Thunb. (2n = 40), *I. makinoi* ‘Hara’ (2n = 40), *I. leucoclada* (2n = 40), and *I. yunnanensis* var. *gentilis* Franch. (2n = 160) (Geukens et al. 2023). Conversely, chromosome number varied greatly in four genera: *i*) *Polygonatum anhuiense* D. C. Zhang et J. Z. Shao (2n = 24), *Polygonatum langyaense* D. C. Zhang et J. Z. Shao (2n = 18), *Polygonatum odoratum* (Mill.) Druce (2n = 18), *Polygonatum zanlanscianense* Pamp. (2n = 28), *P. cyrtonema* (2n = 22) (Sun 1996, Chen and Zhou 2005); *ii*) *Zanthoxylum acanthopodium* Candelle (2n = 64), *Zanthoxylum dimorphophyllum* Hemsley (2n = 36/68), *Zanthoxylum scandens* Blume (2n = 68), *Zanthoxylum oxyphyllum* Edgeworth (2n = 72), *Zanthoxylum tomentellum* J.D. Hance (2n = 72), *Zanthoxylum simulans* Hance (2n =∼132, *Z. nitidum* (2n = 68), *Z. armatum* (2n = 66/98/128/132/136), *Z. bungeanum* (2n = 132/136) (Zhang and Hartley 2008, Chen et al. 2009, Yu et al. 2010, Luo et al. 2018d, Luo et al. 2022b, He et al. 2023, Hu et al. 2023); *iii*) *Bletilla formosana* (2n = 32/36), *B. striata* (2n = 32/34/36/48/51/64/76), *B. ochracea* (2n = 34/36) (Miduno 1954, He et al. 2022c, Huan et al. 2022, Yang et al. 2023); *iv*) *J. regia* and *J. sigillata* (2n = 34) (Luo and Chen 2020), *Juglans* (2n = 32) (Woodworth 1930, Hans 1970, Tulecke et al. 1988, Mu and Xi 1988, Mu et al. 1990). In this study, *P. cyrtonema* (2n = 18), *Z. bungeanum* (2n = 76/134/136/160), *Z. armatum* (2n = 96/100/102/104/132), *Z. nitidum* (2n = 66), *B. formosana, B. striata*, *B. ochracea*, *J. regia,* and *J. sigillata* (all 2n = 34). These results contradict previous studies. The possible causes of inconsistency may be *i*) improper chromosome count in the small and high chromosomes, *ii*) root lignification limiting their chromosome preparation, *iii*) hybridization between closely related species, *iv*) natural or artificial polyploidization, and *v*) apomixis (polyembryo).

Intraspecific chromosome number variation, even as a population, has also been found in species such as *Cuscuta epithymum* (L.) L. and *Cuscuta planiflora* Ten. (García and Castroviejo, 2003). In this case, most variation is attributable to auto- or allopolyploidy. The additional numbers can be explained by ascending or descending dysploidy. Thus, the accumulation of repetitive DNA can lead to an increase in chromosomes and, consequently, to an increase in genome size, especially in subgenus Monogynella (Ibiapino et al. 2022). In the present study, chromosome numbers varied in interspecific and intraspecific populations of the genus *Zanthoxylum*. The cause of the variation is probably similar to that of *Cuscuta* and Monogynella. Furthermore, the stable differentiation in the 5S rDNA FISH pattern between the subgenera suggests that chromosomal rearrangements played a role in splitting the two subgenera. Rather than major structural changes, transpositional events are likely responsible for the variable rDNA distribution patterns among species of the same subgenus with conserved karyotypes (Cai et al. 2006). *Zanthoxylum* genomes have complex chromosome rearrangements, such as chromosomal fission, reversal, and translocations (Hu et al. 2023). This result also further rationalizes the chromosome number variation in this genus. Chromosome polymorphisms within species in natural populations of vertebrates are far less common and are believed to be temporary transitions during chromosomal evolution (Damas et al. 2021, 2022). Likewise, the *Zanthoxylum* may be experiencing chromosomal evolution.

In this study of 64 plants evaluated, the longest chromosome length of each plant was 1.23–6.88 μm, while the shortest chromosome length of each plant was 0.63–3.85 μm, exhibiting strikingly different among the examined species. Previous research accumulated chromosome lengths of hundreds of plant species (Luo et al. 2022a, b, He et al. 2022a, b, d, Luo and He 2021, Liu and Luo 2019, Luo and Chen 2019, Luo et al. 2018d, Luo et al. 2017, Xing et al. 1989, Liu and Sheng 2011). By analyzing these data, it is not difficult to find that chromosome length was slightly different, even for the same accession of the same species. For example, *R. pseudoacacia* 1.12–1.74 μm (Luo et al. 2022b), 0.94–1.67 μm (He et al. 2022a). Nonetheless, chromosome length in the former two literatures was small for both chromosomes (<3 μm). Hence, chromosome length was more suitable for qualitative than for quantitative analyses. Thirty-seven plant species analyzed (more than half) had chromosome lengths lower than 3 μm in this study, consequently dividing them into the small chromosome rank. Because of the hazy centromere mark and tiny chromosomes in many plants investigated, the chromosome size was determined by metaphase and the measurement method. Still, a more delicate karyotype analysis (e.g., arm length, karyotype, and cytotype) was unavailable and limited.

### Occurrence and Diversity of 5S rDNA in Plants

5S rDNA, which occurs in all cellular life forms, is a highly stable tandem repeat sequence that ubiquitously exists in plants (Said et al. 2018). With the evolution and development of the plant, 5S rDNA also underwent simultaneous changes. The length of 5S rDNA in the NCBI database Nucleotide of NCBI was 48–854 bp (Turner et al. 2005, Liu et al. 2017), while its length as a FISH probe in the PubMed database of NCBI was 41–1193 bp (Luo et al. 2017, Islam-Faridi et al. 2020, Glugoski et al. 2020). This study is the first time 5S rDNA testing has been analyzed for 19 species from 13 families. Overall, 5S rDNA occurred in at least two chromosomes in all 64 plants. With advances in science and technology, the occurrence of 5S rDNA has been confirmed in an increasing amount of species (de Barros et al. 2023, de Moraes et al. 2023, Kroupin et al. 2023, Rodríguez-González et al. 2023). Whether the reported length of 5S rDNA is a complete or partial sequence, there is no doubt there is a big difference among these 5S rDNA, including the length and base pair (Röser et al. 2001, Kulak et al. 2002, Moraes et al. 2022).

Because of this, it is quite reasonable that the chromosomally diverse distribution of 5S rDNA is visualized by FISH. Numerous previous studies showed that the numbers of 5S rDNA FISH signal sites ranged from 1 to 71 (Ali et al. 2005, Lan and Albert 2011, Luo et al. 2017, Kovács et al. 2023, Rodríguez-González et al. 2023). Their signal position was found in the chromosome interstitial position, distal position, proximal position, and far away from the chromosome (Amarasinghe and Carlson 1988, Cai et al. 2006, Campomayor et al. 2021, Wang et al. 2022, Rodríguez-González et al. 2023). In this study, 5S rDNA was rather diverse and abundant in signal site number (2–18), position (e.g., interstitial, distal, proximal position, occasionally, outside chromosome), and even as intense (e.g., strong, weak, slight). This is consistent with previous studies of the 5S rDNA signal pattern of *A. fruticose*, *B. formosana* ‘Leshan’, *C. campanulatus*, *H. mutabilis*, *P. nepalensis*, *S. oblata*, two species of *Berberis*, two varieties of *J. regia*, two varieties of *J. sigillata*, two species of *Ligustrum*, two types of *Robinia*, and two varieties of *Z. armatum*, wild/cultural *H. rhamnoides* ssp. *sinensis*, three varieties of *H. rhamnoides*, four species of *Taxus*, and six types of *B. striata* (Luo et al. 2017, Luo et al. 2018d, Liu and Luo 2019, Luo and Liu 2019, Luo and Chen 2019, 2020, Luo et al. 2022a, b). On the contrary, it was different from the 5S rDNA signal pattern of *F. pennsylvanica*, *R. pseudoacacia*, *S. japonicum*, *T. wallichiana* var. *mairei*, *Z. bungeanum* ‘Hanyuanhuajiao’, two types of *B. ochracea*, and two varieties of *Z. armatum* (Luo and Liu 2019, He et al. 2022c, Huan et al. 2022, He et al. 2023) from previous studies. The possible causes of the 5S rDNA signal pattern discrepancy are *i*) lost satellite chromosome with signal and *ii*) variation in different batches of materials in the same species (i.e., intraspecific variation).

### Potential Origin of 5S rDNA Diversity in Plants

Due to the diversity of signal patterns, 5S rDNA was used as an excellent and dynamic marker in the species of *F. pennsylvanica*, *Iris versicolor* L., *L.* − *vicaryi*, *L. lucidum*, *P. nepalensis* (*P. concolor*, former name in Flora of China), *R. pseudoacacia*, *S. oblata* (Lim et al. 2007, Luo et al. 2017, Luo and Liu 2019, He et al. 2022a). After comparing the 5S rDNA in previous studies and in ours, both perfectly reflect the extensive diversity in *P. nepalensis*, which can distinguish all 18 chromosomes according to the signal position, signal intensity, and signal number. Nevertheless, 5S rDNA was quite conserved and dormant in numerous species, such as *C. campanulatus*, *C. sativa*, *H. rhamnoides*, *J. regia*, *M. atropurpureum*, *P. stratiotes*, *P. trichocarpa*, and *Sanguisorba* L. (Mishima et al. 2002, Luo and Chen 2019, 2020, Xin et al. 2020, Alexandrov et al. 2022, Luo et al. 2022a, Stepanenko et al. 2022, de Barros et al. 2023). Why is there such a big difference in 5S rDNA among the above species? There are several plausible hypotheses under investigation, such as *i*) chromosome rearrangement (e.g., deletion, duplication, inversion, translocation), *ii)* polyploidization, *iii*) self-incompatibility, and *iv*) chromosome satellites.

The 5S rDNA position is a hot issue for chromosomal realignment because of its system into long reaches of the standpat tandem repetition unit and its active transcription. This feature implies they are impressionable to chromosomal destruction or non-allelic homologous recombination, thus raising the feasibility of chromosomal realignments, such as fissions, inversions, and fusions (Rosa et al. 2012; Barros et al. 2017; Potapova and Gerton 2019; Warmerdam and Wolthuis 2019; Deon et al. 2020). The 5S rDNA diversification was regarded as variable genomic areas, compliant with double-strand break and chromosomal realignment, facilitating karyotypic reconstruction (Glugoski et al., 2018; Deon et al., 2020, 2022). The 5S rDNA position was also diversified by translocation or transposition events of repeats in those chromosomes (Venancio Neto et al. 2022). An interstitial 5S rDNA position with a diverse location is current, presumably because of a thin inversion. A plus 5S rDNA likely implies the presence of a replication (Coluccia et al. 2020). The 5S rDNA site of one parent was either excluded from the chromosome or shifted into gene silencing and then disappeared, which could decrease the 5S rDNA site number (de Melo and Guerra 2003, Volkov et al. 2017). These studies demonstrate that chromosome rearrangement causes variation in 5S rDNA diversity.

The 5S rDNA has a polyploidization-relevant preference to the distal from a proximal position but keeps a stable loci number (Zhang et al. 2016). The 5S rDNA sites are largely proximal, a highly transparent direction in chromosomes with a single site (Garcia et al. 2016). Consequently, the presence of the 5S rDNA site in the distal chromosomes and the abundance of microsatellites in adjacent areas provide friendly conditions for adding realignments. These results emphasize the effect of variable chromosomal 5S rDNA loci in generating assignments (Gkugoski et al. 2022). No associations between the number of 5S loci and chromosome number, but correspondence with ploidy level and genome size (Adams et al. 2000, Lan and Albert 2011, Garcia et al. 2016), but disputes still occurred (Hasterok et al. 2005, Said et al. 2018, Chai et al. 2022), such as the genus *Cuscuta* L. (García et al., 2017). These studies demonstrate that polyploidization causes variation in 5S rDNA diversity.

### Conclusion

This study conducted FISH-based chromosomal mapping of 5S rDNA markers to provide valid karyotype landmarks to reveal the chromosome number and 5S rDNA signal pattern distribution in 64 plants. Furthermore, we established chromosome physical mapping of each species. Finally, we discuss the proposed origin of 5S rDNA diversity in plants. We are devoted to developing universal oligosequence markers (GAA)_6_, (TTG)_6_, (ACT)_6_, 45S, ITS, and combinatory analysis of additional plant species, particularly woody plants. Altogether, the results reported here enhance the assumption that cytogenetic characteristics (conventional and molecular) could be regarded as excellent markers for chromosome distinction and the presentation and profile of the existing biodiversity in woody plants.

## Data availability

The authors affirm that all data necessary for confirming the conclusions of the article are present within the article, figures, and supplemental figures. Supplemental material available at GENETICS online.

## Acknowledgments

We thank Zhou Yonghong for laboratory equipment support.

